# Localized ROS Generation with UV light in Differentiating Human Neural Progenitor Cells

**DOI:** 10.64898/2025.12.02.691867

**Authors:** Sylvester J. Gates, Wolfgang Losert

## Abstract

Reactive Oxygen Species (ROS) and the associated condition of excess ROS, oxidative stress, has been implicated in a number of diseases including neurodegeneration. However, ROS are also crucial second messengers with beneficial impacts. Within neural cells, ROS signals are known to impact maturation of cells as well as memory and learning. Photobiomodulation (PBM), the use of light to impact cells/tissue, is a promising way to noninvasively modulate ROS. This study investigates the effects of PBM using 370 nm light to increase ROS levels in human neural progenitor cells (hNPC), and study potential impacts on calcium dynamics. We find 370 nm light to be effective at inducing ROS within hNPC. The photoinduction of ROS only impacts ROS levels in illuminated cells, with no measurable signal relay to non-illuminated cells within the acute time period we examined. The increase in ROS generated by our UV light exposure creates elevated basal levels of calcium, but does not impact spontaneous calcium signaling in networks of hNPC cells.

## Introduction

Reactive Oxygen Species (ROS) are reactive molecules and chemicals that have oxygen either as radicals or as non-radicals due to its partial reduction [1–3]. ROS have a dual aspect in biological systems, serving as a both an essential component in certain processes when tightly regulated and being harmful when deregulated or elevated which can cause conditions like oxidative stress and other disease states [4]. In vitro there is therefore a tightly regulated balance between ROS production and antioxidant defense mechanisms which are key to maintaining proper physiological functioning.

ROS are generated as part of general cellular metabolism and are important for several biological functions. A large majority of intracellular ROS are generated within the mitochondria, where they are produced as part of the electron transport chain during oxidative phosphorylation [4–6]. ROS can also be generated within the cytosol, endoplasmic reticulum, peroxisomes, nucleus, andthe plasma membrane [7]. In terms of cell signaling ROS act as secondary messengers in various signaling pathways, regulating processes such as cell proliferation, differentiation, and apoptosis [1, 3, 7].

Elevations in ROS can occur due to external conditions such as radiation, food / alcohol, cigarette smoke, and pharmacological drugs [7]. Within a cell, elevated levels of ROS can cause lipid peroxidation, enzymatic damage & deactivation, as well as DNA damage or mutations [ref]. These cellular changes in turn can cause pathological phenotypes, including DNA damage-induced cancer [ref], neurodegenerative disorders, cardiovascular diseases, and diabetes [7–9]. Since a large increase in ROS levels can be harmful, antioxidants act as a defense mechanism by reducing ROS levels. Antioxidants, which can be molecules, enzymes, or compounds, remove free radicals, thus reducing the amount of ROS in the cell. At the cellular level, key antioxidant enzymes include superoxide dismutase, catalase, glutathione peroxidase, glutathione, as well as originally exogenous compounds like vitamin C & E, carotenoids, and polyphenols [7–9].

In neural cells, ROS plays a crucial role in the context of development, proper functioning, and certain pathological conditions [10]. During neural development from stem cells, ROS acts as a key regulator of proliferation and the downstream differentiation into maturing neuronal cells [11–14]. Once differentiated, neuronal cells still require and use ROS as a key regulator of axonal growth, neurite outgrowth, and polarity [15–18]. In the nervous system, ROS are involved in essential processes such as synaptic plasticity, which underlies learning and memory. ROS also acts as signaling molecules in the regulation of neurotransmitter release, and synaptic maturation and transmission. Drosophila models have shown ROS impacts activity-dependent plasticity and synaptic growth [19,20]. Other research has shown that ROS are implicated in processes required for learning such as long term potentiation (LTP) and GABA-mediated inhibitory neurotransmission [21, 22]. However, within the neural context, excessive ROS (oxidative stress) has been associated with neurodegenerative diseases such as Alzheimer’s and Parkinson’s [1, 19, 23, 24]. Furthermore, ROS-induced DNA damage can cause mutations that can contribute to the pathogenesis of these neurodegenerative diseases. The antioxidant defense systems of the brain are critical in maintaining a delicate balance between ROS generation and elimination, to ensure the proper functioning of neural cells and neurological health [25–28].

Due to the important role ROS plays in both cellular health and disease, photobiomodulation (PBM) stands out as a promising approach to non-invasively modulate the levels of ROS and allow researchers to influence biological processes using light. Photobiomodulation is the use of light to generate biological impacts that can be measured and used to interact non-invasively with cells & tissue [29]. ROS has been seen as a potential target for photobiomodulation both to due its numerous and varied impacts on numerous pathways within cells, and due to the ability to increase ROS levels with near-UV & UV light. Within the literature various cells types have been exposed to near UV light for photobiomodulation: Osteosarcoma [30], Keratinocytes [31], Retinal cells [32]. This suggests that this is a well convserved and ubiquitous process/phenomena.

In this study, we present a non-invasive photobiomodulation approach to induce increased ROS levels in developing living human neural network progenitor cells (hNPC), and investigate how the duration and power of UV illumination affects ROS generation. Using precisely patterned UV light fields on an in vitro network of hNPC cells, we investigate how increases in ROS impact neighboring cells not directly exposed to UV light, and how the altered ROS levels impact calcium signaling

## Materials and Methods

### Culturing and maintenance of hNPC

Human neural progenitor cells (hNPCs) were derived from human fetal brain tissue of the ventral mesencephalon (Millipore Sigma, #SCC008), which includes the v-myc transgene. The cells were cultured following previously established methods. For expansion, cells were plated on Geltrex [Get geltrex manufacturer] coated tissue culture flasks or plates (NEST and Corning) and maintained in a growth medium consisting of DMEM:F12 + GlutaMax (Thermo Fisher Scientific), supplemented with 2% B-27 Plus Neural Cell Supplement (Thermo Fisher Scientific), 1% penicillin-streptomycin (Sigma-Aldrich), 50 *µ* M heparin (Stemcell Technologies), 10 ng/mL basic fibroblast growth factor (bFGF, Stemcell Technologies), and 20 ng/mL epidermal growth factor (EGF, Stem-cell Technologies). For passaging, cells were detached using Trypsin ([Get Brand]) for 2-5 minutes. After detachment, one volume of DMEM:F12 + GlutaMax was added to collect cells, and then cell solutions were centrifuged at 300 g for 5 minutes. Resulting cell pellet was resuspended in fresh growth medium and replated on prepared Geltrex-coated plates. For differentiation, growth medium was modified by excluding the growth factors (bFGF and EGF). Cells were cultured in a humidified incubator at 37°C with 5% CO2 and were passaged at approximately 95% confluence. All experiments were conducted using cells between passages 5 and 20.

### Intracellular ROS Visualization

Intracellular reactive oxygen species were visualized using CellROX Deep Red Reagent (Thermo Fisher Scientific), following the manufacturer’s protocol. A 5 mM stock solution of CellROX Deep Red (CRDR) was prepared, and an appropriate dilution was made to achieve a final working concentration of 1 µM in the cell culture medium. Cells were incubated with the reagent for 60 minutes at 37°C. After incubation, cell samples were rinsed with PBS, and medium was replaced with prewarmed imaging solution. Imaging was performed immediately using a widefield fluorescent microscope system equipped with temperature, humidity, and CO_2_ control. To visualize The fluorescent indicator CRDR samples were excited using 640 nm light.

### Intracellular Calcium Labeling & Visualization

Intracellular calcium was labeled using Calbryte-590 AM (AAT Bioquest) according to the manufacturer’s protocol. A stock solution was prepared in anhydrous DMSO. To achieve a final working concentration of 1 µM, the appropriate dilution of Calbryte-590 AM was added to the cell culture medium. Cells were incubated with the labeling solution for 30 to 60 minutes at 37°C. After incubation, the medium was replaced with a prewarmed imaging solution, which could be either the cell culture growth medium Imaging was performed immediately on a system equipped with temperature, humidity, and CO_2_ control. The fluorescent indicator was excited using a 561 nm laser.

### Imaging/Microscopy

All imaging experiments were performed using a Ti-2 Eclipse microscope. All images were captured using a 20x water-immersion objective lens for wide-field imaging. Imaging excitation was provided using a 648 nm laser for CellRox Deep Red for fluorescent imaging of ROS or using 560 nm laser for the Calbryte 590 for fluorescent imaging of calcium. Stimulation was performed using the microscope’s DMD system (Polygon Mightex 1000) to stimulate using 370 nm LED. Fluorescence was detected using a Teladyne Kinetix CMOS camera. Image acquisition was controlled via NIS-Elements software. Cells were imaged under a controlled environment at 37°C with 5% CO2 live-cell incubation system.

### Image & Statistical Analysis

All image analysis was performed using custom Python scripts. Image sequences from movies with dwell time and intensity modulations were loaded in. Each movie FOV was segmented using a static image based on where stimulation occurred, as well as a single rectangular region of comparable size as a control/background region. To generate z-score normalized, blurred difference image, z score normalized frames, one from the beginning and one from the end, of stimulation were subtracted from one another followed by the application of a Gaussian blur.

On a field-of-view basis each stimulated region of interest’s average intensity over time was collected and first normalized to the first frame starting level via subtraction. The background signal was then subtracted from the normalized stimulation. Finally, each FOV was normalized per replicate basis so that at the end of the 99%, 3 minute stimulation was 1 - this ensured that every value in each FOV was scaled based on its starting value and where we expect the intensity to be greatest (highest percentage and dwell time of stimulation). All replicates were then combined to find the mean normalized intensity over time. Linear interpolation was performed on the first 3 frames was used to calculate the slopes for each condition based on the LED stimulation intensity. One-way analysis of variance (ANOVA) performed to analyze slopes at various LED intensity levels showed a significant p-value. To determine differences between groups, Tukey’s HSD was performed for the 33, 66, and 99% LED intensities. P-value for significance was set at 0.5.

To measure local response/impact of LED stimulation on ROS, around a particular ROI an expanded 30 pixel dilation was performed successively to define regions of interest. Python was used to calculate the average normalized intensity in each ROI around the 3 minute stimulation for all LED intensities tested. The slope was estimated via linear interpolation as described above. To determine significance, the average initial slope was used with an applied Bonferroni correction applied, comparing against each successive region of interest within each LED intensity tested.

To process calcium data fluorescence time-lapse data is organized to include calcium data from both before stimulation and after stimulation within a single field of view. For each pair, cellpose is run on the final frame of the before-stimulation movie to segment the image into cellular regions. These cellpose masks are then further split into categories based on whether the cell was in the stimulated region or in a non-stimulated region. The median relative change is calculated for every individual cell of the last frame before-stimulation and the first frame after-stimulation and stored as a per-cell metric. To quantify the response of cells to stimulation based solely on their calcium level, the mean relative change and the standard deviation are calculated based on the non-stimulated cells, and a threshold set at two standard deviations. The proportion of cells in both the non-stimulated and stimulated regions is calculated. To determine if the threshold proportion of the stimulated cells is significantly different from the non-stimulated cells a a paired T-test is performed with a p-value set at 0.05.

## Results

### 370 nm Light Can Generate Reproducible Levels of ROS in developing hNPC

To determine whether hNPC are susceptible to photoinduced ROS generation via 370 nm light and how controlled this generation is, developing hNPC (post differentiation initiation day 7-9) were exposed to 3 power levels and 3 dwell times in nine defined regions within a single field of view (Fig. 1A). We chose power levels of 33, 66, and 99% DMD illumination power, as well as dwell times of 20 seconds, 60 seconds, and 3 minutes. Various light intensity and dwell times were applied in a single field of view over the course of a single recording (Fig. 1B). The regions of interest (ROI) were exposed as follows: 1) 20 s, 33%, 2) 20 s, 66%, 3) 20 s, 99%, 4) 1 min, 33%, 5) 1 min, 66%, 6) 1 min, 99%, 7) 3 min, 33%, 8) 3 min, 66%, 9) 3 min, 99%. Each ROI were illuminated at separate yet consecutive time points during the recordings (Fig. 1B). In a single ROI, when exposed to 370 nm light we could determine a change in fluorescent intensity by segmenting the region and measuring the average signal, for visualization (Fig. 1C).

**Figure 1:**
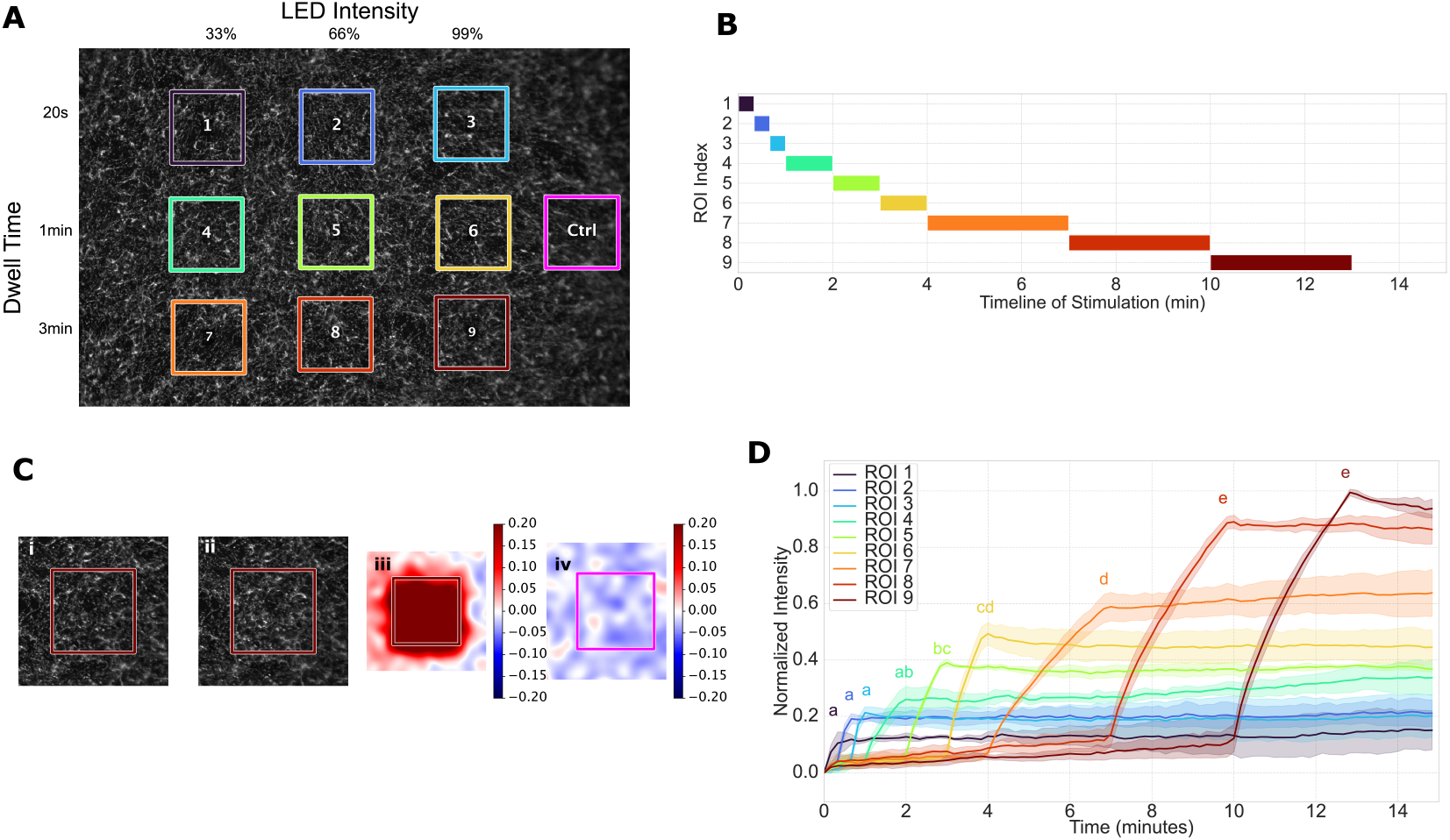
Experimental Setup and Response of 370 nm Photoinduced induced ROS in hNPC. Representation of a single experimental field of view showing various colored regions of interest to be stimulated with various LED intensity and dwell time (A). Timeline of stimulation events separated by ROI index; color matches that found in A (B). Representative example from ROI 9 showing frames before and after stimulation (i & ii respectively), along with z-score normalized difference image of the subsequent frames (iii) from ROI 9, and the control region(iv)(C). Mean normalized intensity by region (shown in colors) across the entire experimental run time (n=3 biological replicates)(D). Letters represent subgroups that do not contain significantly different change in intensity during stimulation.

We analyzed the change in fluorescent intensity across each of our 9 ROI to understand how both power and dwell time affected ROS generation measured via the ROS indicator. The normalized intensity for each ROI was calculated by first normalizing the intensity time course to the initial frame. Next, a dynamic background (derived from a normalized non-stimulated background ROI) was subtracted from all ROIs. Finally, the corrected intensity was normalized per biological replicate to the maximum intensity observed across all ROIs at a specific reference point (end of final stimulation, frame 78), setting this maximum value to 1. This normalized intensity quantification showed that increasing both dwell time and laser power over the tested time led to further increases in fluorescent intensity measured via the ROS indicator (Fig. 1D). Applying analysis of variance (ANOVA), we found a statistically significant difference in change in intensity during stimulation based on individual ROIs (F=129.085, p < 0.001). A Tukey’s post-hoc test for pairwise differences found 5 distinct statistically significant subsets (a,b,c,d,e)(p < 0.05). ROI 1-4 belong to the non-significant group a. ROI 4&5 belong to group b. ROI 5&6 belong to group c. ROI 6&7 belong to group d. And ROI 8&9 belong to group e. Generally, there was a positive trend showing that ROI with longer duration of stimulation (corresponding to higher index number) or higher LED intensity were more likely to have a statistically significant greater change in intensity change during stimulation (compared to those with a smaller ROI index number).

### 370 nm generated ROS in hNPC is dose dependent

To look at the potential dose-dependent nature of 370 nm photoinduced ROS in hNPC, we aligned the change in intensity from the group-specific stimulation start point of our LED intensities tested and found each appeared to have specific kinetics based on the LED intensity used for stimulation (Fig.2A). To quantify this change, we looked at the initial slope during the first 20 seconds of stimulation for the ROIs with 1- and 3-minute stimulation (means: 33% = 0.0049 ± 0.0005, 66% = 0.0074 ± 0.0006, 99% = 0.0097 ± 0.0011). A one-way analysis of variance was performed with a significant p-value (*p* < 0.0001) that indicates a significant difference among the group means. Tukey’s HSD was performed in a pairwise manner with results indicating all initial slopes were significantly different from one another (33% vs 66%, *p*_adjusted_ = 0.0002; 33% vs 99%, *p*_adjusted_ = 0.0; 66% vs 99%, *p*_adjusted_ = 0.0001) (Fig.2B).To determine dose-dependence we initially measured the normalized change in fluorophore intensity from the beginning to the end of stimulation for the 9 stimulated ROIs. A Coherent FieldMaxII-TO Laser Power Meter (1098579) power meter was used to measure LED intensity of the full DMD and then converted into estimate measurements based on the area of stimulation. These measurements were then used to determine the total energy dose in mJ applied to our 9 stimulated ROIs. We constructed a dose-dependence curve and categorized each of our ROIs based on the LED intensity used and then applied a linear fit for each LED intensity group (Fig.2C).

**Figure 2:**
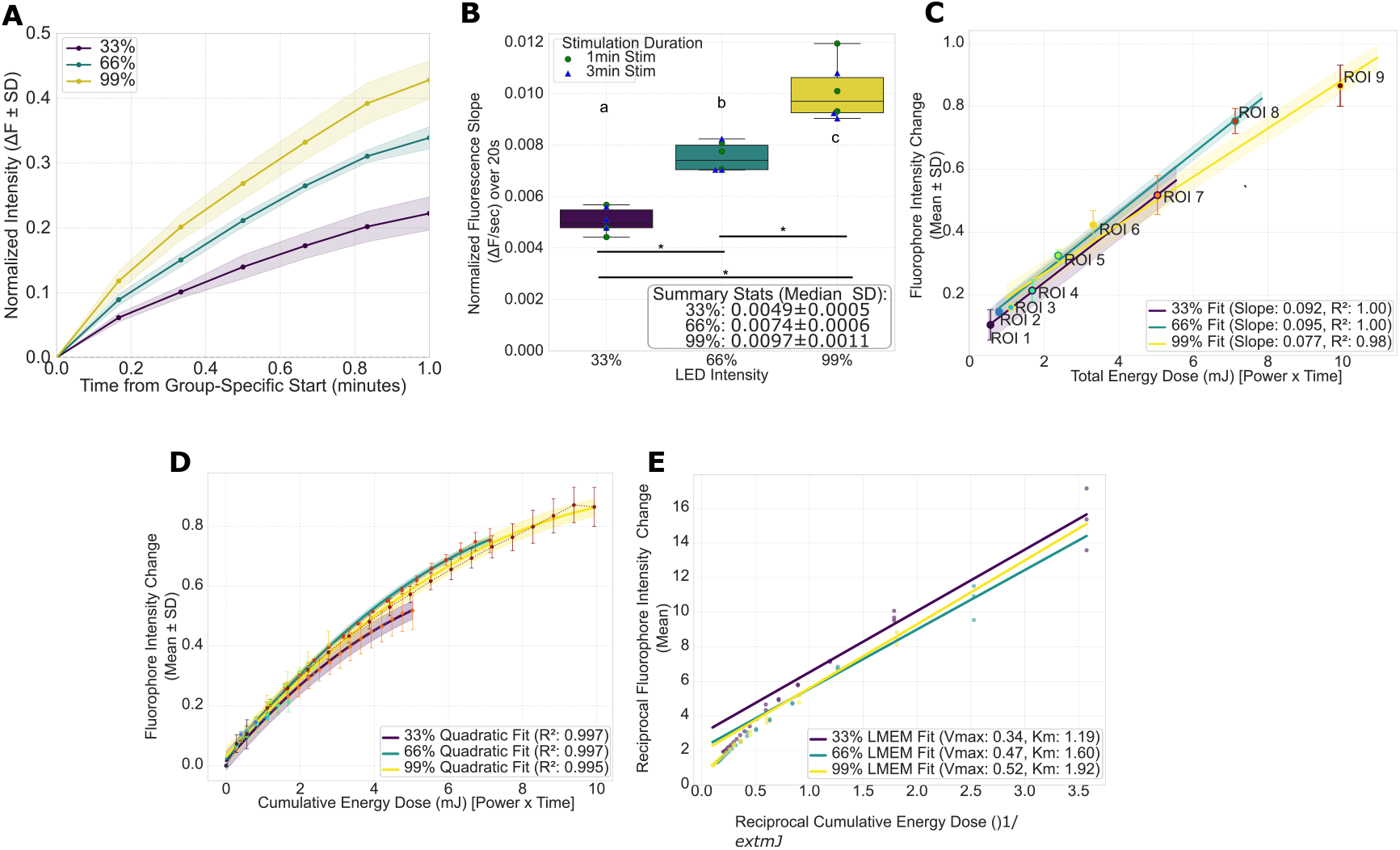
370 nm Photoinduced induces ROS in hNPC in dosage dependent manner. Mean normalized fluorescence intensity over time grouped by intensity, and aligned to stimulation onset, shaded regions represent standard deviation across biological replicates (n=6) (A). Box plots for slope of fluorescence intensity during the first 20 seconds of stimulation grouped by LED intensity (B). Groups with different letters represent conditions with significantly different means. Dose-Response curves of fluorophore intensity change vs total energy dose based on stimulated ROIs with linear fit based on LED intensity (C). Error bars represent SD for each ROI, shaded bands represent SD for linear fit. Dose-response curve for continuous data for each ROI during the duration of stimulated time with quadratic fit based on LED intensity (D). Error bars represent SD for each ROI, shaded bands represent SD for quadratic fit. Lineweaver-Burk plot of double reciprocal data for reciprocal intensity change vs reciprocal energy dose with LMEM fit based on LED intensity (E).

To determine if different LED intensity groups have an impact on the ROS indicator intensity change when accounting for energy dose, we first performed Homogeneity of Regression Slopes (HoRS) to check that relationship between LED intensity group and our slopes. The interaction term for LED intensity was not statistically significant (F = 1.56, p = 0.343) indicating the dose-response relationship (slope) across the LED intensities did not differ significantly. With the assumption of HoRS satisfied, a standard one-way Analysis of Covariance (ANCOVA) was performed to test the effect of LED intensity on the change in intensity, controlling for dose (mJ). We found dose was highly significant (F = 257.51, p < 0.001) as a strong predictor of change in intensity. Whereas LED intensity group was not significant (F = 0.873, p = 0.473). We therefore concluded that there is no difference in the adjusted mean change in fluorophore intensity when controlling for dose across the three LED intensity groups tested. This evidence suggests that total energy dose is the primary factor driving the change in ROS indicator fluorophore intensity - this suggests ROS accumulation in hNPC due to 370 nm light is a dose-dependent response.

To further confirm this, as opposed to using the final change in the fluorescent intensity from start to end of stimulation, we calculated the change in fluorescent intensity from the start to all other time points during stimulation. This allowed us a continuous variable that we could measure for each of our 9 stimulated ROIs which we used to create a dose-response curve (Fig.2D). When we combined LED intensity groups, we could fit a quadratic across each (Fig.2D). However, these data were not suitable for ANCOVA due to multiple, repeated, continuous measurements across each of our spatially independent ROI samples, which appear to show saturation. Based on the dose response curve we believe the relationship between intensity change and cumulative energy dose may follow a saturation-based, Michaelis-Menten-like non-linear model. We used a linear-mixed effects model (LMEM) proxy using double reciprocal transformation of the data to model the impact of LED intensity group on saturation parameters. LED intensity was modeled as a fixed effect, while group membership was random. A Lineweaver-Burk plot was generated based on the LMEM analysis for visual validation of the linearized saturation model (Fig.2E) LMEM was performed using restricted maximum likelihood method with the reciprocal change as the dependent variable (N = 78 observations, 9 groups, log-likelihood = -46.7101). The LMEM successfully converged with the intercept (the 33% LED intensity group) coefficient being 2.982. Our model found a highly significant relationship with reciprocal dose (p < 0.001), but neither LED intensity group nor interaction between LED group and dose were significant (p > 0.05 in all four cases). Most of the variation between groups was due to random intercepts (*σ*^2^ = 3.105) due to between-group variability in baseline signals. Altogether, this further suggests that reciprocal change in intensity as a function of reciprocal dose is statistically equivalent across the three LED intensity groups tested. This leads to the conclusion that there is a dose-dependent response of intensity change based on cumulative energy dose, when accounting for changes in each individual ROI over time.

### Photo-induced ROS is locally constant

To assess whether ROS generated locally may impact or lead to cell-cell signaling or the spread of ROS, we measured the levels of ROS in five successive, 10 micron-wide regions surrounding the stimulated regions (Fig. 3A&B). We chose to focus on the 3 minute stimulation dwell times as that provided the most data points to better understand if regions surrounding the stimulated region showed similar metric changes based on the increase in fluorescent intensity (Fig. 3B). From the normalized intensity curves we determined the initial slope, the change in intensity during stimulation, and the area under the curve (AUC). For each of these metrics we then constructed graphs to visualize the change based on both LED intensity as well as relative ROI number as a proxy of distance from the stimulated target region (Fig. 3C-E).

**Figure 3:**
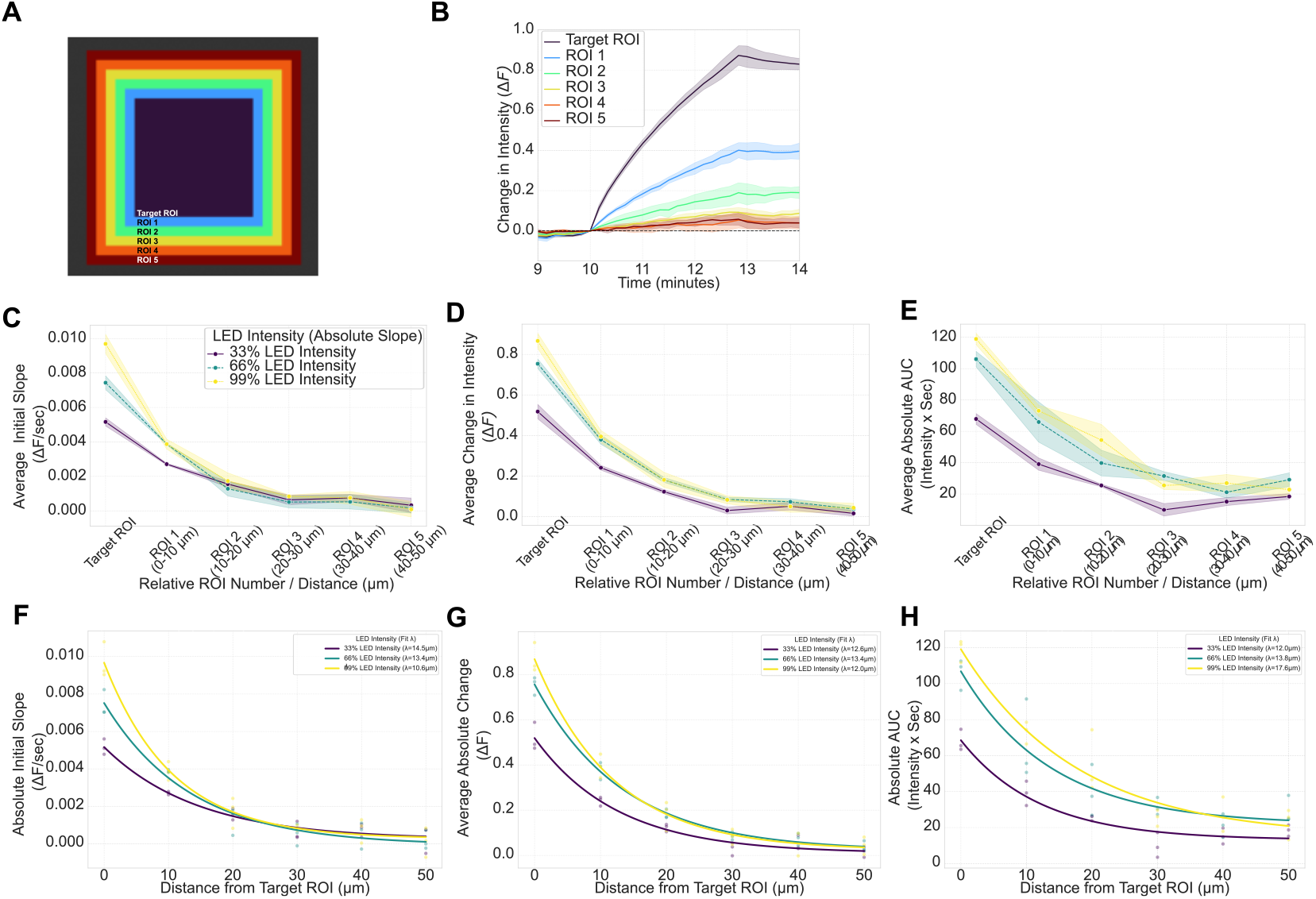
370 nm Photo-induced ROS remains localized within stimulation region with exponentialdecay. Schematic representation of a targeted stimulation region (same as in Figure 1) with successive 10 µm regions around it (regions 1-5) shown in various color (A). An example for the 99% LED intensity, 3 min stimulation during experimental time, with stimulation starting at the 10 minute mark, aligned y=0 at the start of stimulation; shaded regions represent standard deviation of biological replicates (B). Average initial slope during the first 20 seconds (C), average change in intensity during stimulation (D), and average AUC during entire stimulation (E) for the target region and successive stimulation regions; shaded regions represent standard deviation of biological replicates. Exponential modeled curves for initial slope (F), absolute change during stimulation (G), and AUC during stimulation (H) over distance in microns of the tested stimulation intensities. *λ* for each metric in legend.

To test the effects of LED intensity and the relative ROI number (as a proxy for distance) on the fluorescence intensity metrics of initial slope, change in intensity during stimulation, and area under the curve – a two-way analysis of variance (ANOVA) was performed. The ANOVA revealed statistically significant main effects for both LED intensity and relative ROI number for all three metrics explored. For initial slope, LED intensity (F(2,36) = 44.63, p < 0.0001) and relative ROI number (F(5,36) = 29.25, p < 0.0001) were highly significant. For change in fluorescent intensity during stimulation, LED intensity (F(2,36) = 60.02, p < 0.0001) and relative ROI were both highly significant (F(5,36) = 70.37, p < 0.0001). And for AUC LED intensity (F(2,36) = 23.48, p < 0.0001) and relative ROI (F(5,36) = 15.30, p < 0.0001) were again, both highly significant. These indicate that the spatial pattern is dependent on the intensity setting for all three metrics.

To determine where the differences were we we performed Tukey’s HSD post-hoc test to perform all pairwise comparisons across our treatments groups. When we compare the low baseline signal (33% LED intensity at ROI 5) to other sources we find a consistent trends between metrics and within metrics. For initial slope we found that for all LED intensities, ROI 1 was the furthest ROI we could find a significant difference compared to the low baseline signal. For absolute change in intensity we found for the 33% LED intensity, ROI 1 was the furthest away ROI to have a significant difference, whereas for 66% and 99%, ROI 2 was the furthest away ROI with a difference compared to the low baseline signal. For AUC we find that for 33% LED intensity only the stimulated region is significantly different, however for 66%, ROI 1 was the furthest away ROI to have a significant difference, while for 99%, ROI 2 was the furthest away ROI to have a significant difference compared to the low baseline signal. In this analysis the change in fluorescence is the most sensitive to detecting difference compared to the lowest signal (33% LED intensity at ROI 5) baseline, as it provides the most statistically different results. However, AUC appears useful for detection limit as it shows the clearest trend and delineation of the LED intensities based on the difference compared to the low signal baseline. Altogether these analyses show that only within about 10 *µ* does initial slope of all LED intensities show significant differences compared to low intensity baseline. For higher LED intensities (66 and 99) the change in fluorescence is different from baseline at up to 20 *µ* m, while 33% is only 10 *µ* m compared to baseline. And finally, AUC appears the most interesting, with significant differences in AUC being seen further out based on LED intensity. With significant difference from low signal baseline being see only in the stimulated target region for 33% intensity, at only 10 *µ*m distance for 66% intensity, and at up to 20 *µ*m for 99% intensity (Fig. S4).

We modeled this interaction using an exponential decay model to determine the space constant to determine how far the response spread based on our LED intensity (Fig.3F-H). Although the spatial constant *λ* was not significant for any of the metrics found (ANOVA p = 0.7677, 0.7750, 0.4647 for initial slope, change in intensity, AUC respectively), our model did find trends in *λ*. For the initial slope *λ* showed an inverse relationship with LED intensity (14.5 → 13.4 → 10.6*µm*) suggesting fastest spread occurs only near stimulation. The change in fluorescent intensity during stimulation showed very little change based on LED intensity (ranging from 12.01 to 13.43 *µm*). Finally, we found a direct relationship between AUC and increasing LED intensity (12.0 → 13.8 → 17.6*µm*) which may suggest higher LED intensity leads to higher spread of total signal response (AUC).

### Photo-induced ROS in hNPC increases Calcium concentration locally

To determine whether ROS photobiomodulation within hNPC could be used to alter cellular dynamics, we measured the impacts of 370 nm light stimulation on calcium dynamics in hNPC. Cells were treated with both the ROS indicator (CRDR) as well as a calcium indicator (Calbryte590). After incubation cells were first imaged briefly for calcium dynamics for 2 minutes, next, a single CRDR ROS indicator image was captured, then a square target region within the field of view was stimulated using 370 nm light (3 minutes, at 99% power), after which a single CRDR ROS indicator image captured, and finally followed by 2 more minutes of calcium imaging (Fig. 4A). As expected cells stimulated with 370 nm light showed a corresponding increase in ROS (Fig. 4B). We also found that the calcium level of cells stimulated in a square target region had an increased florescent intensity compared to cells that were left untreated within the same field of view (Fig. 4C). To determine if this change was significant, we segmented cells and looked at the change in intensity on a per cell basis both before and after stimulation. We found that a higher proportion of stimulated cells were above a threshold value of intensity (2 sigma) of the non-stimulated cells - suggesting higher calcium intensity (Fig. 4D). We found that the stimulated cells had a significant higher proportion of cells exhibiting mean intensity greater than 2 sigma (p = 0.0209). Next, to analyze changes in calcium florescence intensity over time in of our cells due to localized stimulation, we took pre- and post-stimulation image sequences and determined statistical measures of pixel intensity over time including the variance, standard deviation, the sum of differences between all frames, and root mean square. Next we calculated the average of each metric in our stimulated and surrounding non-stimulated region of interest. To normalize we then calculated the ratio of the average value in image sequence before stimulation to the image sequence after stimulation for each ROI for each metric. We found that only the standard deviation was significantly different for our stimulated cells (p = 0.024) while the normalized variance, sum of differences, and root mean square did not differ significantly (p=0.053, 0.074, and 0.052 respectively). (Fig.4D)

**Figure 4:**
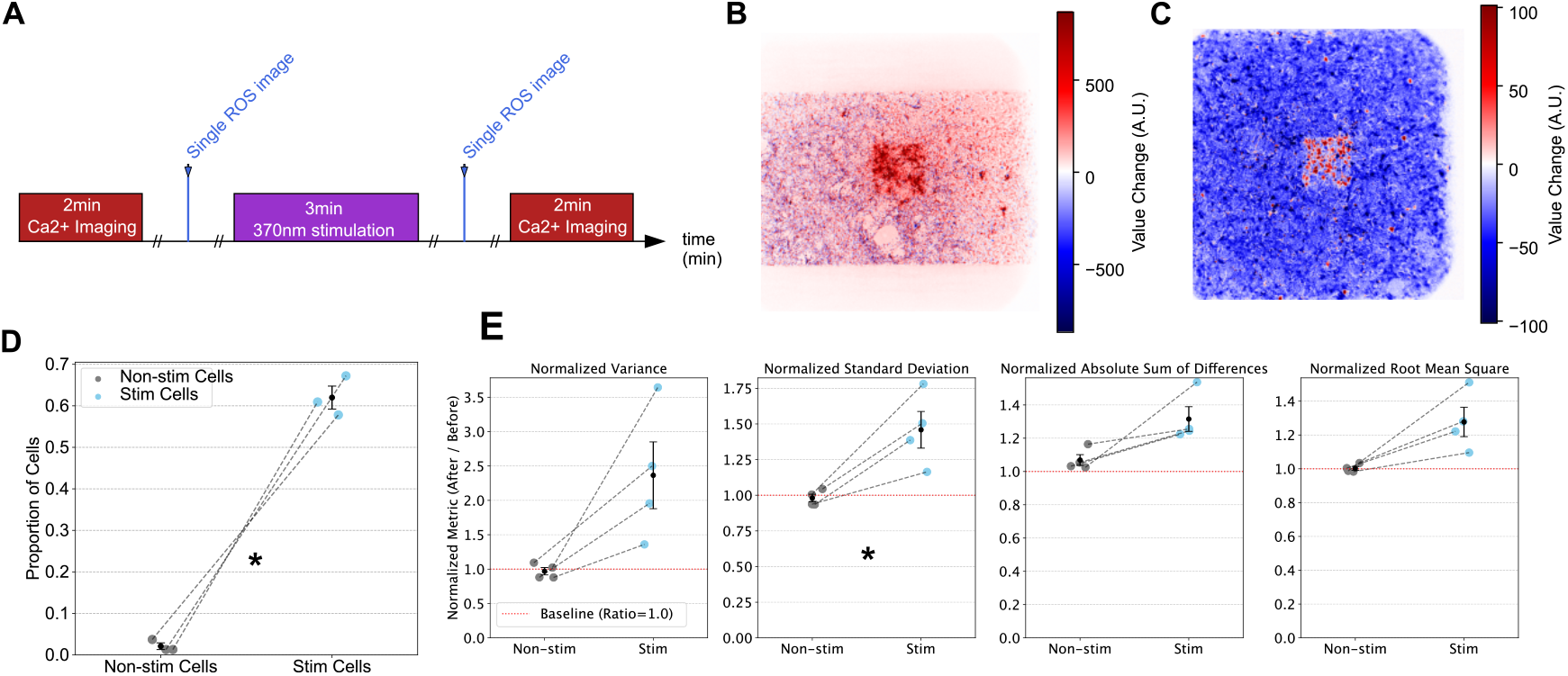
370 nm Photo-induced ROS increases basal local calcium level. Schematic of imaging paradigm for calcium imaging, ROS imaging, and 370 nm stimulation (A). Processed image from a single FOV which shows relative change in fluorescent intensity of ROS indicator from the difference of the frames before and after stimulation (B). Processed image from a single FOV which shows the relative change in fluorescent intensity of calcium indicator from the difference of the frames before and after stimulation; note the bright central square region which is where stimulation was applied (C). Proportion of cells that are above 2 standard deviations of intensity from the non-stimulated cells mean intensity for both non-stimulated and stimulated cells of frames before and after stimulation (n=4 biological replicates)(D). Metrics quantifying normalized variance, standard deviation, absolute sum of difference, and root mean square respectively for non-stimulated and stimulated cells for entire movies of dynamics (n=4) (E).

## Discussion

In this study, we employed a network of hNPC cells to characterize how photobiomodulation may impact ROS levels and calcium dynamics within a neural network. The use of developing hNPCs allows us to model neural networks consisting of mixtures of astrocytes, neurons and other cell types. Using a DMD to localize light and with live cell imaging, we have shown that 370 nm light can be used as a photobiomodulation tool to noninvasively probe and modulate hNPCs via ROS generation.

First, we have shown that 370 nm light exposure of hNPCs induces ROS, which is assayed via a fluorescent ROS indicator (Fig. 1). UV light exposure generates excess ROS immediately upon exposure, with increasing ROS levels with exposure duration and intensity, as expected. We show that 370 nm induced increases in ROS follows a dose-dependent response using both statistical analysis of the raw data and modeling (Fig.2). While we cannot determine the absolute values of ROS, we note that the increase in ROS levels remains spatially localized even after the stimulation ended (Fig.3). We also show that the local increase even outside of the stimulated region appears to change AUC based on LED intensity - which might suggest scattering. The lack of change increases in intensity after stimulation has subsided is consistent with an absence of cellular stress signals relayed between cells. This could indicate that UV exposure modulates ROS in a range small enough to avoid triggering strong secondary cellular stress responses. Nevertheless, we find that photoinduced ROS has a measurable impact on some cellular signaling: we find elevated calcium levels within hNPC networks stimulated with UV light, but again the increased calcium levels are localized to the region of the neural network exposed to UV light (Fig.4). The UV light power is a limiting factor in our study, and it is possible that higher amounts of ROS generated via 370 nm light may also alter the dynamics of calcium in hNPCs. Another limitation with this current study was that we only looked at dynamics within our experimental range (less than 15 minutes). Future work should look at the impacts of 370 nm light in hNPC over longer periods of time to determine if PBM has other impacts at later time points.

Overall, this study characterized the powerful regulatory capacity of spatially patterned UV light. Our findings indicate that near UV light exposure yields powerful control over neural networks, allowing us to alter both metabolic factors (ROS) and signaling molecules (calcium) precisely, with cell scale spatial control and second scale timing control. The rapid change of ROS levels is particularly useful given our recent finding on in-vitro primary rat hippocampal neural networks, that collective spike timing-dependent plasticity is notable within minutes [33]. Thus, in future studies, UV triggered modulations of ROS and calcium levels may be used for precise interventions to alter information processing and learning of living neural networks.

## Funding Information

Funding was provided by the AFOSR Biophysics Program under grant FA9550-25-1-0002

## Disclosures

## Acknowledgments

We acknowledge and thank Dr. Michael Denton and AFRL for fruitful discussions. We also would like to thank Dr. Kan Cao for discussions and initial contributions of the hNPC.

## Notes

### Competing Interest Statement

The authors have declared no competing interest.

