## Supplementary Figures for "Localized ROS Generation with UV light in Differentiating Human Neural Progenitor Cells"

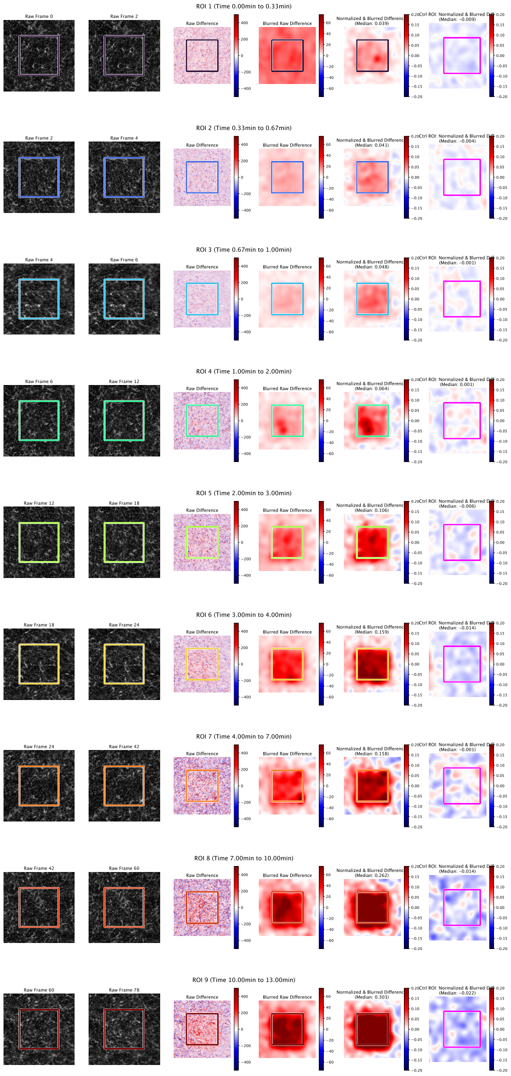


**Figure S1: Processed Difference Image From Single Field of View**

Each row represents one of the 9 ROIs stimulated from a single field of view in an experiment. Columns are as follow: (1) first frame of stimulation, (2) last frame of stimulation, (3) raw difference image between first and last frame of stimulation, (4) gaussian blurred raw difference image, (5) z-score normalized and gaussian blurred difference image, (6) control region with performed z-score normalized and gaussian blurred difference image.

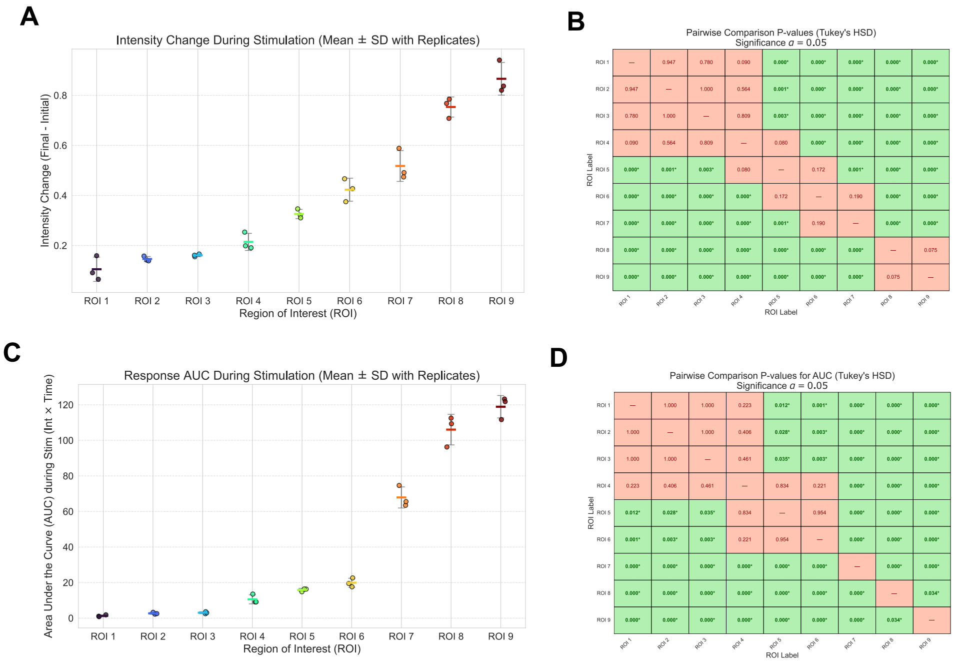


**Figure S2: Pairwise Comparisons of Change in Intensity and AUC during stimulation**

Bar plots showing change in normalized fluorescent intensity from beginning to end of stimulation per ROI; error bars represent standard deviation (A). Heatmap representing Tukey's HSD pairwise comparison p-values of data shown in A (B). Bar plots showing AUC of normalized fluorescent intensity from during stimulation per ROI; error bars represent standard deviation (C) Heatmap representing Tukey's HSD pairwise comparison p-values of data shown in C (D).

A

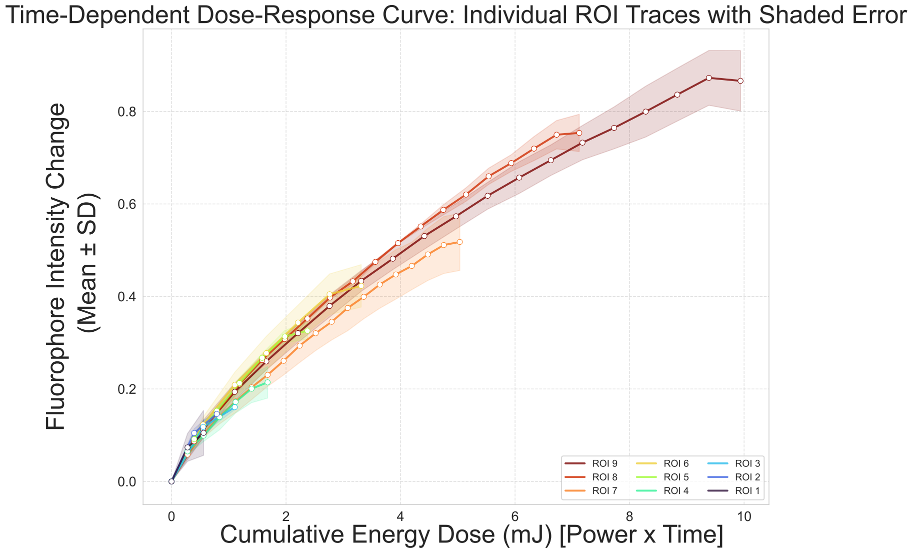


**Figure S3: Time-Dependent change in fluorophore intensity based on energy dose per ROI**

Fluorophore intensity change vs cumulative energy dose for all ROIs studied. Continuous data for each are from successive time points throughout stimulation. Shaded error represents standard deviation of the data (N=3).

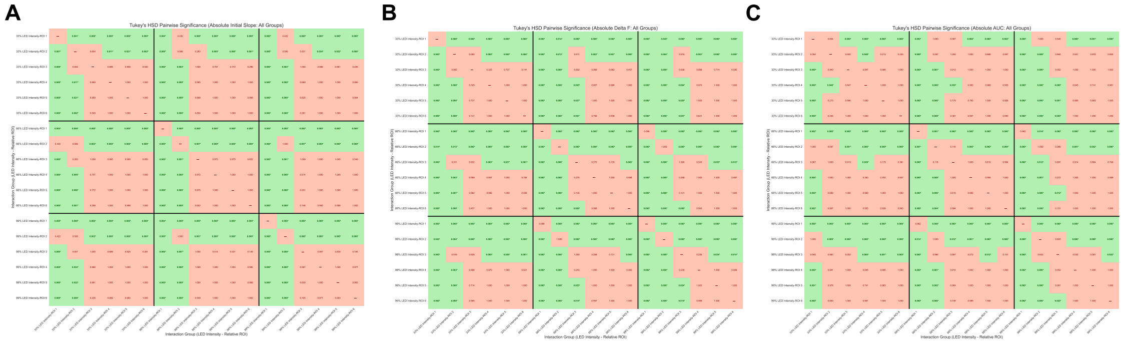


**Figure S4: Tukey's HSD Heatmap of Pairwise Comparison P-Values for LED Intensities Over Successive Distances**

Initial Slope (A). Change in normalized flourescent intnesity during stimulation (B). AUC over stimulation (C).
